# Predicting dysfunctional age-related task activations from resting-state network alterations

**DOI:** 10.1101/678086

**Authors:** Ravi D. Mill, Brian A. Gordon, David A. Balota, Jeffrey M. Zacks, Michael W. Cole

**Affiliations:** Center for Molecular and Behavioral Neuroscience, Rutgers University, Newark, NJ, 07102, USA; Department of Radiology, Washington University in St Louis, St Louis, MO, 63110, USA; Department of Psychological and Brain Sciences, Washington University in St Louis, St Louis, MO, 63130, USA

**Keywords:** functional connectivity, task activation, fMRI, Alzheimer’s, aging

## Abstract

Alzheimer’s disease (AD) is linked to changes in fMRI task activations and fMRI resting-state functional connectivity (restFC), which can emerge early in the timecourse of illness. Study of these fMRI correlates of unhealthy aging has been conducted in largely separate subfields. Taking inspiration from neural network simulations, we propose a unifying mechanism wherein restFC network alterations associated with Alzheimer’s disease disrupt the ability for activations to flow between brain regions, leading to aberrant task activations. We apply this activity flow modeling framework in a large sample of clinically unimpaired older adults, which was segregated into healthy (low-risk) and at-risk subgroups based on established imaging (positron emission tomography amyloid) and genetic (apolipoprotein) risk factors for AD. We identified healthy task activations in individuals at low risk for AD, and then by estimating activity flow using at-risk AD restFC data we were able to predict the altered at-risk AD task activations. Thus, modeling the flow of healthy activations over at-risk AD connectivity effectively transformed the healthy aged activations into unhealthy aged activations. These results provide evidence that activity flow over altered intrinsic functional connections may act as a mechanism underlying Alzheimer’s-related dysfunction, even in very early stages of the illness. Beyond these mechanistic insights linking restFC with cognitive task activations, this approach has potential clinical utility as it enables prediction of task activations and associated cognitive dysfunction in individuals without requiring them to perform in-scanner cognitive tasks.

**Significance Statement:** Developing analytic approaches that can reliably predict features of Alzheimer’s disease is a major goal for cognitive and clinical neuroscience, with particular emphasis on identifying such diagnostic features early in the timeline of disease. We demonstrate the utility of an activity flow modeling approach, which predicts fMRI cognitive task activations in subjects identified as at-risk for Alzheimer’s disease. The approach makes activation predictions by transforming a healthy aged activation template via the at-risk subjects’ individual pattern of fMRI resting-state functional connectivity (restFC). The observed prediction accuracy supports activity flow as a mechanism linking age-related alterations in restFC and task activations, thereby providing a theoretical basis for incorporating restFC into imaging biomarker and personalized medicine interventions.

## Introduction

Analyzing the synchronization of activity fluctuations in task-free rest is thought to provide insight into the brain’s intrinsic network organization (Raichle, 2010; Petersen and Sporns, 2015). Formalized in the subfield of resting-state functional connectivity (restFC), such approaches have primarily been applied to human functional magnetic resonance imaging (fMRI) data. As testament to the reproducibility of restFC, convergent network topologies have been recovered across different regional atlases (Power et al., 2011; Ji et al., 2019), non-fMRI human imaging modalities (Brookes et al., 2011; Kucyi et al., 2018) and non-human animals (Wang et al., 2013; Stafford et al., 2014). Whilst these findings highlight that restFC is reliably observed, debate persists over its cognitive relevance, given that the experimentally unconstrained nature of rest raises practical difficulties in separating signal from noise (Power et al., 2012; Laumann et al., 2016), as well as theoretical difficulties in moving from an exploratory to explanatory understanding of brain network function (Mill et al., 2017).

Evidence of restFC’s cognitive relevance comes from clinical research linking restFC alterations to pathology (Buckner et al., 2008; Buckholtz and Meyer-Lindenberg, 2012). Given the severe prognosis and societal burden of Alzheimer’s disease (AD), much of this work has interrogated restFC changes characterizing various forms of healthy and unhealthy aging (Andrews-Hanna et al., 2007; Sorg et al., 2007; Ferreira and Busatto, 2013; Geerligs et al., 2015; Ferreira et al., 2016). Burgeoning research focuses on restFC changes in early at-risk stages of AD (Hedden et al., 2009; Sheline and Raichle, 2013; Schultz et al., 2017), as neurobiological abnormalities such as elevated positron emission tomography (PET) measures of amyloid beta may precede clinical impairment by many years (Jack et al., 2010; Sperling et al., 2011). Alzheimer’s-related alterations of restFC emerge around the same time as elevated amyloid (Sheline and Raichle, 2013) and have been associated with presence of the genetic apolipoprotein (APOE) *ε*4 allele (Sheline et al., 2010), suggesting utility of restFC in developing imaging biomarkers to expedite diagnosis and intervention.

Recent reports have extended towards using restFC to quantitatively predict or classify age-related conditions (Dosenbach et al., 2010; Woo et al., 2017; Du et al., 2018). However, failures of predictive models generalizing out-of-sample (Onoda et al., 2017; Teipel et al., 2017; Fountain-Zaragoza et al., 2019) highlight limitations of entirely data-driven approaches to predicting Alzheimer’s-related pathologies, especially as artifactual contaminants of restFC can drive clinical group differences (Siegel et al., 2016; Hodgson et al., 2017). These findings again call for increased efforts to clarify the cognitive relevance of restFC to identify clear mechanisms by which restFC alterations impact on AD and other pathologies.

To this end, we recently developed an approach inspired by neural network simulations to mechanistically predict task activations from restFC via the concept of activity flow (the propagation of task-evoked activity between neural populations; Cole et al., 2016; Ito et al., 2017). This approach directly interrogates the cognitive relevance of restFC, given that tasks are designed to elicit particular cognitive processes. Furthermore, in addition to restFC, fMRI task activations are disrupted in various forms of aging (Grady, 2012; Campbell and Schacter, 2017), raising the potential for age-related alterations of restFC and task activations to arise from a common activity flow mechanism.

The present report sought empirical evidence for this mechanism (Figure 1). We hypothesized that unhealthy age-related restFC alterations disrupt the ability for activity to flow between brain regions, leading to the emergence of dysfunctional task activations. We tested this framework in clinically normal older adults, a subset of whom presented positively for PET amyloid or APOE risk factors for AD, which were separately used to define ‘at-risk’ subjects. We found that activity flow mapping reliably predicted task activations in held-out at-risk AD subjects from their pattern of restFC. This result supports activity flow alterations as a mechanism underlying age-related dysfunction, even in the earliest stages of AD.

**Figure 1.**
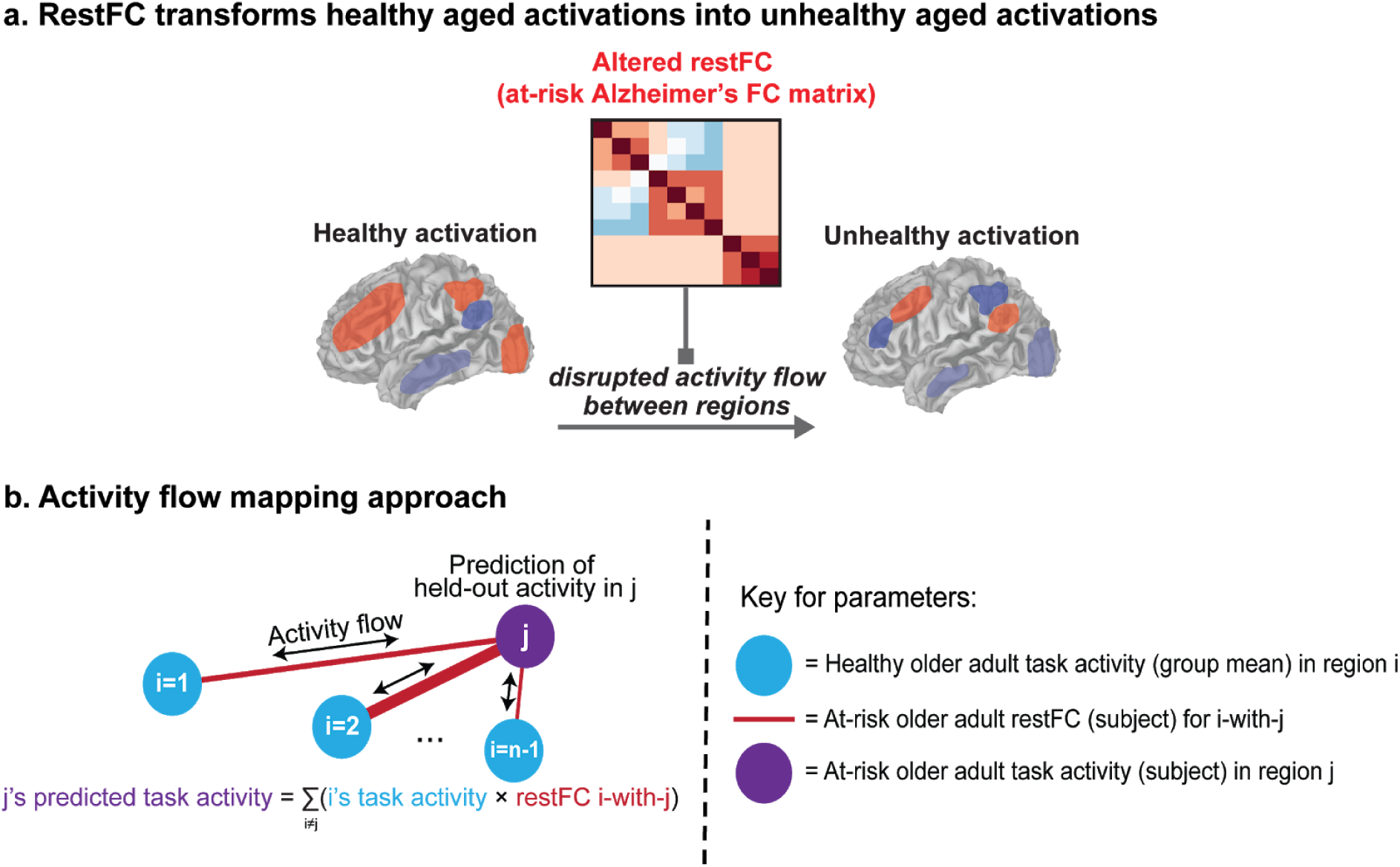
Theoretical and methodological principles underlying application of activity flow mapping to build an integrated connectivity-activity account of at-risk Alzheimer’s disease (AD). **a)** Abstract theoretical underpinnings of activity flow mapping – which is based on the core mechanism underlying neural network simulations – in a clinical context. A ‘healthy’ aged task activation pattern is transformed into an ‘unhealthy’ aged activation (i.e. one associated with being ‘at-risk’ for developing AD) by an altered pattern of resting-state functional connectivity (restFC) in at-risk subjects disrupting the ability for task activations to flow between brain regions. **b)** Methodological formalization of the activity flow mapping approach. Task activation in a held-out brain region (j) in a held-out at-risk AD subject is predicted on the basis of the dot product between the activation state in the rest of the brain (regions i, estimated as the mean activation in the healthy group) and the restFC between i and j (estimated from the to-be-predicted at-risk subject). See Method for further details.

## Materials and Methods

### Participants

The sample comprised 101 right-handed elderly subjects (age mean=65.1 years, age range=42-82 years, 63 female) collected as part of the Adult Children Study at the Knight Alzheimer’s Disease Research Center at Washington University in St Louis (Knight ADRC, https://knightadrc.wustl.edu/). Full details of behavioral and neuroimaging measures acquired by the Knight ADRC project are available in Gordon et al (Gordon et al., 2015). Subjects included in our sample underwent behavioral (Mini Mental State Exam, MMSE; Folstein et al., 1975) and neuropsychological (Clinical Dementia Rating, CDR; Morris, 1993) assessment, rest and task fMRI scans, structural MRI scans, ^11^[C] Pittsburgh Compound B (PiB)-PET imaging for high levels of amyloid beta uptake, and a DNA swab testing for presence of the apolipoprotein (APOE) *ε*4 allele. All subjects were assessed as clinically ‘normal’ at the time of recruitment (Mini Mental State Exam, MMSE score > 24; Clinical Dementia Rating, CDR=0). All subjects passed the exclusion criteria of high motion across runs (average absolute movement >1.50 mm or an average relative movement of 0.5 mm), presence of neurologic damage (stroke or traumatic brain injury), and a lag between their fMRI and PiB-PET imaging sessions greater than 90 days.

### Segregation of sample into at-risk and healthy aged groups

We binarized the APOE *ε*4 and PiB-PET amyloid beta measures separately to provide distinct methods of segregating our sample into ‘at-risk AD’ (i.e. ‘unhealthy’) and ‘healthy’ subject groups. This enabled us to test the generalizability of the activity flow mapping approach across different ways of identifying at-risk subjects. The APOE segregation was created by labeling any subject with at least one APOE *ε*4 allele as at-risk, and all other subjects as healthy (see Table 1 for demographic information). At-risk subjects were identified in the amyloid segregation on the basis of a standardized PiB uptake ratio (SUVR) greater than 1.42 (following Jack et al., 2017; Vlassenko et al., 2018; see next section for further details). Note that our definition of at-risk AD encompasses both previously adopted at-risk (APOE *ε*4 genotype) and preclinical (elevated PET amyloid) AD categorizations. For brevity, the Results section focuses primarily on the APOE segregation due to the larger at-risk group obtained in comparison to the amyloid segregation (see Table 1). However, the pattern of observed results is virtually identical for both segregation types (see final section of Results).

**Table 1.**
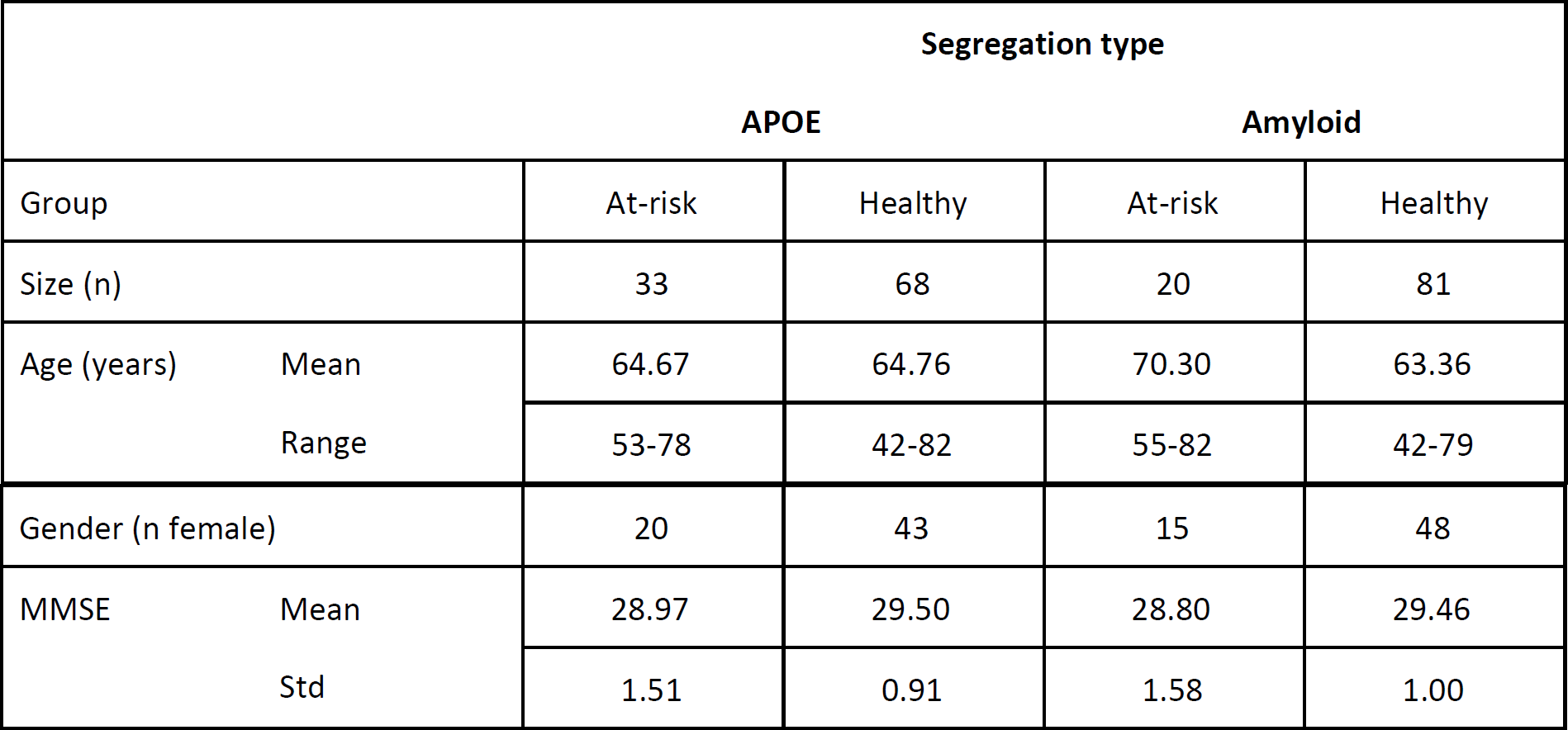
Demographic information for the two at-risk AD segregation approaches: APOE and amyloid. MMSE=Mini Mental State Exam scores; Std=standard deviation.

### Data acquisition and preprocessing

MRI data was collected on a Siemens Trio 3T scanner. Anatomical T1-weighted images were acquired via a magnetization-prepared rapid gradient-echo sequence (MPRAGE; TR=2400ms, TE=3.16ms, flip angle=8°, field of view=256mm, 1mm isotropic voxels, sagittal orientation). Task fMRI images were acquired using an interleaved whole-brain echo planar imaging (EPI) sequence (TR=2000ms, TE=25ms, flip angle=90°, field of view=256mm, 4mm isotropic voxels, 36 slices in interleaved sagittal orientation). Task fMRI data was collected for two tasks (see next section for paradigm details): semantic animacy (2 runs, 303 volumes each) and color Stroop (2 runs, 295 volumes each). Resting-state fMRI images were acquired using a similar EPI sequence (TR=2200ms, TE=27ms, flip angle=90°, field of view=256mm, 4mm isotropic voxels, 36 slices in interleaved sagittal orientation; 2 runs, 164 volumes each). Separate dual-echo gradient-recalled echo (GRE) fieldmaps were also acquired to correct b0 distortions in the task and rest EPI images respectively.

T1 images were segmented using Freesurfer (Fischl, 2004). fMRI preprocessing was conducted in FSL using the FEAT toolbox (Woolrich et al., 2001). Subjects’ task and rest fMRI images underwent motion correction, fieldmap b0 unwarping, slice timing correction, linear coregistration to their anatomical T1 and non-linear normalization to an age-appropriate MNI template (created from a separate large sample of demographically matched older adults; Gordon et al., 2015). Subsequent analyses were conducted at both the regionwise and voxelwise levels. For the regionwise analyses, task and rest timeseries were extracted from 264 regions from the Power functional atlas (Power et al., 2011). For the voxelwise analyses, timeseries were extracted from all gray matter voxels. Nuisance regression was performed for the regionwise/voxelwise timeseries via general linear models (GLMs), which included regressors for 6 motion parameters, white matter and ventricular timeseries, and their temporal derivatives. The GLM fit to the task fMRI data also included regressors for the two tasks (see next section for details). The residual timeseries from the rest GLM and the beta activation amplitudes from the task GLM were used for the main activity flow mapping analyses. The voxelwise beta activation maps were spatially smoothed using non-Gaussian nearest-neighbor averaging (at 4mm), which reduces spatial autocorrelation versus Gaussian smoothing.

Full details about PiB-PET acquisition in the Knight ADRC project was provided previously (Su et al., 2013). Briefly, after injection of the PiB tracer, subjects underwent 60 minute dynamic PET imaging scans using a Siemens 962 HR+ ECAT scanner. A summary measure of whole-brain PET amyloid was estimated from anatomical atlas regions identified from Freesurfer segmentations of each subject’s T1 image (using the ‘wmparc’ segmentation; Fischl, 2004). Errors in the Freesurfer segmentation were identified and corrected manually. The Freesurfer regions of interest were then aligned to the native PET images, and the standardized uptake value ratio (SUVR) was computed as the median PiB uptake at each region relative to a cerebellar reference (following Rousset et al., 2008; Su et al., 2015). SUVR was averaged across regions to provide a summary measure of whole-brain PET amyloid deposition, which was then binarized to label at-risk AD and healthy subjects for the Amyloid segregation (whole-brain SUVR > 1.42=at-risk; Vlassenko et al., 2018).

### Experimental task design and fMRI task activation estimation

The design of the two tasks is detailed in Figure 2a. Each task was briefly practiced by subjects just prior to their fMRI sessions. Both tasks involved word stimuli presented in a block design, with the presentation format and durations matched closely across them. There were 2 runs of each task, each alternating between 4 task blocks (containing 24 trials with jittered intertrial intervals) and 5 rest intervals. The semantic animacy task preceded the color Stroop task, and required subjects to judge whether individually presented words referred to living or nonliving things. Words used in this task were balanced across living/nonliving categories in terms of length, orthographic neighborhood and frequency. The color Stroop task required participants to judge whether words were presented in a red or blue font; the words’ meaning was either ‘congruent’ with the font color (e.g. the word ‘red’ in red font), ‘incongruent’ (e.g. the word ‘blue’ in red font), or ‘unrelated’ to color (e.g. the word ‘deep’ in red font). Stimulus classes for both tasks were balanced within task blocks (e.g. equal numbers of living and non-living words were presented for each animacy task block, in a randomized order).

**Figure 2.**
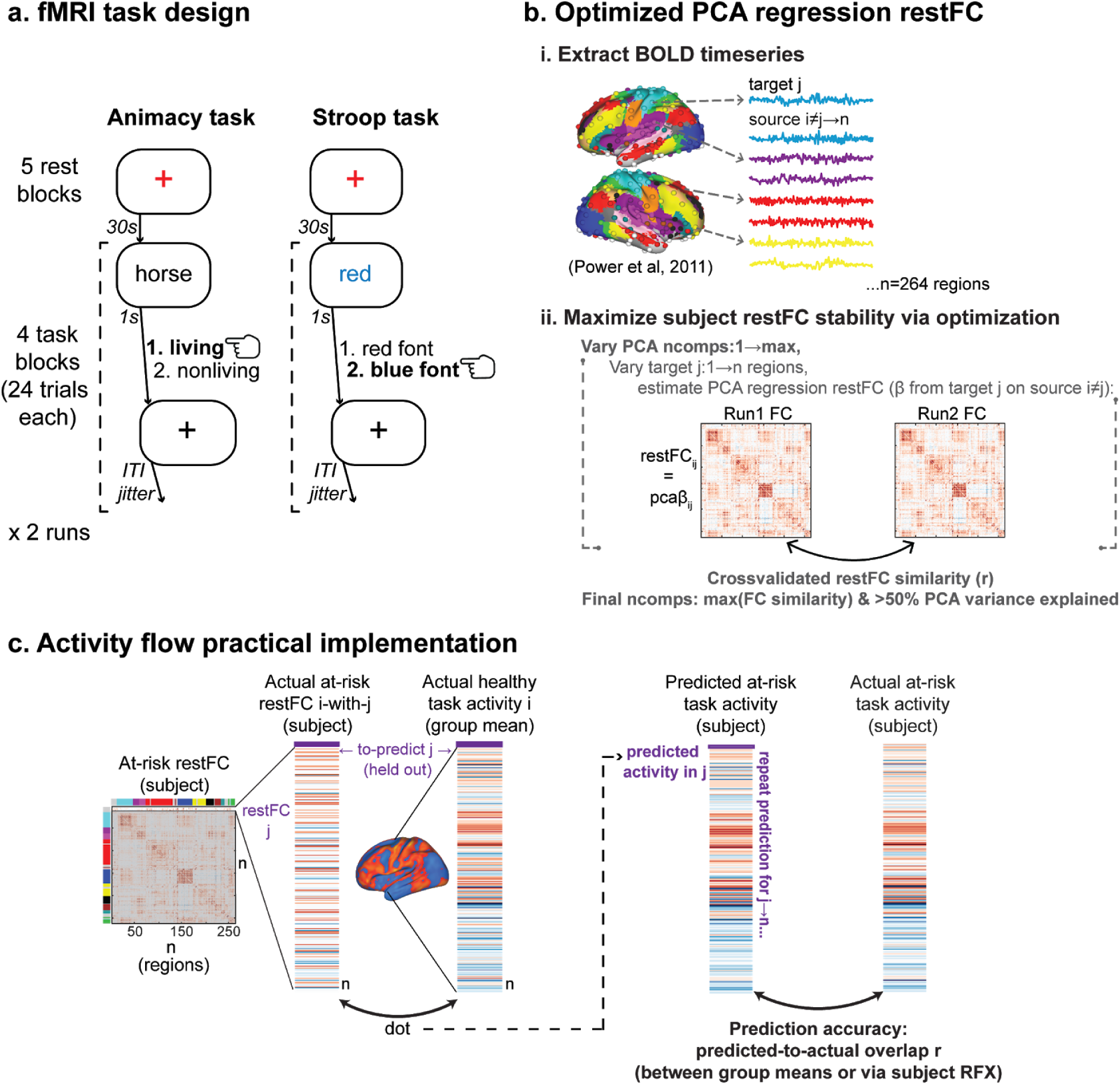
Task design, restFC estimation and activity flow mapping implementation. **a)** Design schematic for the two fMRI tasks: semantic animacy and color Stroop. See Method for a detailed description. Example trials are depicted for each task, with the hand symbol denoting the correct response. ‘ITI jitter’=intertrial interval jitter, which ranged from 1-9s in increments of 2s. **b)** Approach to estimating restFC for each subject using an ‘optimized’ form of multiple linear regression with PCA dimensionality reduction. After extraction of BOLD rest timeseries from functional regions-of-interest (color-coded according to network affiliations from the Power et al atlas, 2011), restFC was estimated for each subject via a regression model predicting target j region timeseries from the principal components of the remaining source i region timeseries. The stability (i.e. matrix similarity) of the restFC solutions across the 2 runs was optimized for each subject via a cross-validation scheme that is described in the Method section. ‘ncomps’=the number of principal components from the PCA of the source i timeseries included in the regression model predicting j. This is the term that is varied as part of the restFC optimization. **c)** Practical implementation of activity flow mapping approach. Note that the restFC term used to generate the activity prediction for a given region/voxel j is a single row from that at-risk AD subject’s restFC matrix (color-coded in the Figure according to the Power network affiliations). ‘subject RFX’=quantification of prediction accuracy via a subject-level random effects approach (see Method for details).

fMRI task activation amplitudes were estimated via GLM in Matlab. In addition to nuisance regressors (see previous section), the task GLM included two regressors modeling blocks for each task as boxcars convolved with the canonical hemodynamic response function (using the ‘spm_hrf’ function from the SPM toolbox). GLMs were fit to the regionwise and voxelwise timeseries data separately, and the resulting beta amplitude estimates were used in all analyses involving fMRI task activations.

### Resting state functional connectivity estimation

The residual timeseries from the rest fMRI GLM were used to estimate restFC. Figure 2b illustrates the approach, which involves an optimized form of multiple linear regression with principal component analysis (PCA) dimensionality reduction. Multiple linear regression has been used for activity flow mapping previously (Cole et al., 2016), with a primary benefit over alternative FC estimation methods being the removal of ‘indirect’ connections. For example, an underlying ground truth FC pattern with connectivity between regions A-B-C would incorrectly yield connections between A-C if using Pearson correlation, but not multiple linear regression. The optimization of multiple linear regression FC was performed for each individual subject, with the goal of maximizing the inter-session stability of restFC and prevent overfitting to noise. The latter may arise from multiple linear regression FC estimation using the maximum number of principal components permitted by the data (i.e. the minimum across timepoints and region variable dimensions), given that some components will reflect noise rather than neural signal. Our optimization approach varied the number of principal components included in the regression in a within-subject cross-validation scheme, which aimed to maximise the similarity of the whole-brain pattern of restFC across the two resting-state runs per subject.

Specifically, for a ‘target’ brain region j, a PCA projection was performed for all ‘source’ regions i→n excluding that target region (see Figure 2b). The number of principal component scores (ncomps) retained after this step was iteratively varied from 1 to the maximum permitted (i.e. 164 timepoints per rest run minus 1 = 163 max ncomps). The retained PCA scores were then used in a multiple linear regression model predicting the target j timeseries, with the resulting coefficients multiplied by the PCA loadings to obtain betas capturing resting-state connectivity between region j and all other regions i (pca *β*_ij_ in Figure 2b). This process was repeated for all target regions j to populate the region x region restFC matrix for each rest run. For each ncomps cross-validation loop, the similarity between the run 1 and run 2 restFC matrices was calculated as the Pearson correlation between all off-diagonal matrix elements. The optimized ncomps value was chosen as that yielding the maximum FC similarity between rest runs, whilst also accounting for greater than 50% of the variance explained in the source region PCA (averaged across all PCAs performed on source regions accompanying each target region loop). These constraints ensured that FC stability was maximized without excluding signal components accounting for meaningful variance in individual rest runs. Hence, each subject had a different optimized ncomps value (mean ncomps across subjects=10.26, std=6.88; mean restFC similarity=.58, std=.09; mean source PCA variance explained=63.90%, std=10.85), and this value was used to estimate multiple regression restFC for the subject’s full timeseries concatenated across runs 1 and 2. Note that this final step avoids circularity given that the optimization was performed to maximize within-subject restFC stability, rather than to directly optimize any analyses that we report inferential statistics for (e.g. the strength of the activity flow mapping predictions).

A separate restFC optimization was performed for the voxelwise analyses following an almost identical approach, wherein target voxels j were predicted by the PCA projection of source voxels i. The only modifications were the exclusion of spatially proximal voxels from the source voxel set (i) i.e. voxels from the same functional region as the target voxel and all voxels within 9mm of that functional region were excluded. Functional region affiliations were identified from the Gordon functional atlas (Gordon et al., 2016), as this parcellation provides regional affiliations for every gray matter voxel in the brain, unlike the Power functional atlas which only provides affiliations for 10mm diameter spheres. This step reduced the inflation of activity flow mapping predictions by spatial autocorrelation of the fMRI BOLD response.

### Machine learning classifications using restFC and task activation features

We trained multivariate machine learning classifiers to distinguish between at-risk and healthy subjects, separately on the basis of regional restFC and regional task activation features. We employed a representational similarity analysis approach (RSA; Kriegeskorte et al., 2008; Diedrichsen and Kriegeskorte, 2017; Hebart and Baker, 2018; Spronk et al., 2018) with leave-one-subject-out cross-validation. To increase the sample size for classifier training, we concatenated at-risk subjects across both APOE and amyloid segregation types, yielding a combined at-risk group size of 40. Given that the healthy group was larger (n=61) than the at-risk one, we selected a random subset of 40 subjects from the healthy group to ensure that training was performed on balanced group sizes.

Features for the restFC classification were the off-diagonal elements from each subject’s restFC matrix (264 × 264 regionwise connections), as estimated via the optimized regression approach described above. Features were first z-scored within subjects. Given the high dimensionality of restFC (69432 connections), we performed a feature selection step at the start of each training/testing loop to confine the classification to the most highly discriminative connections (Dosenbach et al., 2010; Vergun et al., 2013). These connections were identified as those that significantly differed in a 2-sample t-test contrasting at-risk versus healthy group restFC at an alpha threshold of .01. The held-out subject was excluded from these 2-sample t-tests to prevent circularity. Note that use of a standard statistical threshold as a feature selection criterion offsets the risk of overfitting compared to optimization of this parameter in the present dataset (e.g. via nested cross-validation over a range of possible alpha thresholds). After confining to discriminative restFC features, at-risk and healthy group restFC templates were computed by averaging features for each group separately (again, excluding the held-out test subject). Pearson correlation was used as a distance metric to assess the similarity between the test subject and both templates, with a binary classification decision set by comparing whether the test subject’s restFC was more similar to the unhealthy group (yielding an ‘unhealthy’ classification) or healthy group (yielding a ‘healthy’ classification) template. Decisions were averaged across test subjects to estimate classification accuracy, with significance assessed via binomial test against 50% chance classification.

The approach to the machine learning classification using task activations was almost identical to the restFC classification. Given the low dimensionality of the regional activations, we used all 264 activation features without performing a feature selection step. Separate classifications were performed for the two fMRI tasks, as well as after averaging activations across the two tasks for each subject to promote ‘task general’ activation features.

### Activity flow mapping approach

Figure 1b depicts the abstract approach in using activity flow mapping to predict task activations in at-risk AD subjects. Figure 2c provides a more practical guide to implementing the procedure. For a to-be-predicted at-risk subject, task activations in held-out regions (‘j’ in Figure 1b) were predicted by the dot product of the averaged healthy group task activations in the remaining regions (‘i’ in Figure 1b) and the at-risk subject’s restFC between j and i. The restFC estimates thus weight the ability for activity to flow between regions, as indicated by the variable width of the red lines in Figure 1b. Iterating through all brain regions populated a predicted task activation vector for each at-risk AD subject (see Figure 2c). The accuracy of the activity flow predictions was then assessed by computing the overlap (Pearson correlation) between the predicted task activation vector and the actual activation vector. This was done both after averaging predicted and actual activations across subjects (group-level overlap) and for each subject’s individual vectors (subject-level overlap).

Statistical significance of the prediction accuracy was assessed in both parametric and non-parametric fashion. For parametric statistics, the significance of the group-level overlap was indexed by the p value accompanying the Pearson correlation. The subject-level significance was assessed in a random effects approach, with the overlap r values for each subject transformed to Fisher-z values and submitted to a one-sample t-test against 0. We also conducted non-parametric permutation tests to: i) demonstrate the importance of region-to-region correspondence between the healthy group activation and at-risk subject restFC terms to the prediction accuracy, and ii) deal with any violation of the independence assumption in the parametric statistics. The latter is a possibility given that each region’s activation prediction is made on the basis of dot product operations between task activation and restFC terms, which involve overlapping regions that could introduce dependency. Hence for 1000 permutations, the group healthy activation term and the rows of each at-risk subject’s restFC matrix were shuffled. Activity flow predictions were then computed from the dot product of these shuffled terms as described above, ultimately generating permuted group-level and subject-level overlap r values. Significance was assessed by calculating the proportion of permuted r values that were higher than the observed values (group- and subject-level), at an alpha threshold of .05.

Activity flow mapping was applied separately for each task and at-risk AD group segregation type (APOE and amyloid; see Table 1). The procedure was virtually identical for both the regionwise and voxelwise analyses. For the voxelwise analyses, accompanying the exclusion of nearby voxels from the restFC estimation for a given voxel (see earlier Method section), we also excluded the same set of voxels from the group healthy activation vector when computing the activity flow prediction for that voxel. Combined with use of non-Gaussian smoothing, this step prevented inflation of activity flow predictions by fMRI BOLD spatial autocorrelation amongst proximal voxels.

We also applied a ‘two-cycle’ variation of this general approach, wherein the predicted at-risk task activations obtained from the standard ‘one-cycle’ approach described above were multiplied by subjects’ restFC a second time. This was theoretically motivated to enhance the ability for activity flow mapping to transform the healthy activation state to an at-risk AD (unhealthy) activation state (see Figure 1a), and hence improve the overlap of the predictions with the actual at-risk activations (see Results for further details).

To clarify, the task activation term for this activity flow model is taken from the healthy group, whereas the restFC term is taken from each to-be-predicted at-risk AD subject (see Figure 1b and Figure 2c). This represents a modification to the original activity flow mapping approach (Cole et al., 2016), wherein both activation and restFC terms were taken from the to-be-predicted subject. This modification permits application of the procedure in cases where task fMRI data is unavailable or difficult to acquire for a given subject, which raises clinical utility.

### Activity flow prediction of dysfunctional activations: regressing-out approach

The previous section describes our approach to using activity flow mapping to predict task activations in at-risk subjects. We also applied modified versions of the approach to more directly predict dysfunction associated with at-risk AD i.e. differences in task activation between the at-risk and healthy groups. To achieve this, we used a regression approach in which we firstly regressed out the group healthy task activation from the predicted activation for each at-risk subject. After this operation, the residual activation vector reflects differences between the healthy group and that at-risk subject (scaled by the constant in the regression). We also regressed out the group healthy activation from the actual activation for the same at-risk subject. This enabled comparison of the overlap between the predicted and actual dysfunctional activation vectors, estimated via Pearson correlation as before. Statistical significance of the overlap was assessed by both parametric and non-parametric (permutation) tests, applied to the group-level and subject-level overlap. This approach was repeated for both the regionwise and voxelwise data (again, excluding nearby voxels from the activation and restFC terms that generate the voxelwise predictions).

The permutation test approach was similar to that described in the previous section: scrambled group healthy activation and subject at-risk restFC vectors were generated for each to-be-predicted at-risk subject. Activity flow was then computed from these terms, with the same unscrambled group healthy activation vector regressed out of the predicted and actual activations prior to computing their overlap at both the group level (after averaging activations across subjects) and the subject level (for each individual subject). This generated a permuted distribution of activity flow mapping prediction overlap (over 1000 permutations), to which the observed group-/subject-level overlap was compared at an alpha threshold of .05.

### Activity flow prediction of dysfunctional activations: contrast image approach

As an alternative method of predicting dysfunctional task activations, we used activity flow mapping to generate a predicted between-group contrast image that captured activation differences between the at-risk and healthy groups. We focused on the voxelwise data for this analysis as this spatial scale provides the most visually interpretable results, and as this is typically the scale at which between-group contrast images are analyzed in fMRI research. Task activation maps for all healthy and at-risk AD subjects were firstly zscored (within subjects) to more clearly recover the pattern of activation. The average of the healthy subject activations was then subtracted from the average of the activity flow-predicted at-risk subject activations. The resulting ‘predicted contrast image’ directly indexed how activations at each brain voxel differed in the at-risk relative to the healthy group. This predicted contrast image was correlated with the ‘actual contrast image’ (the averaged actual at-risk activations minus the averaged healthy activations) to assess how accurately task activation dysfunction was captured.

To correct for any inflation of the contrast image overlap induced by subtracting the same vector (healthy group activation) from two other vectors (the predicted and actual at-risk group activations) that are subsequently correlated, we used a non-parametric permutation approach to generate a more appropriate null than overlap r=0. Over 1000 permutations, we generated activity flow-predicted task activations based on scrambled group-averaged healthy activation and scrambled at-risk subject restFC terms (as per the permutation tests described in previous sections). From the resulting scrambled predicted group at-risk activation, we subtracted the unscrambled group healthy activation. This permuted predicted contrast image was correlated with the observed (unscrambled) actual contrast image. Significance of the observed overlap values was assessed against this permuted null distribution at an alpha threshold of .05.

We also conducted a thresholded version of the voxelwise contrast image analyses. A 2-sample t-test contrasting the actual at-risk AD activations versus the healthy activations was used to identify the most highly dysfunctional voxels in the actual data. A ‘predicted’ version of this 2-sample t-test was also computed contrasting predicted at-risk activations versus the healthy activations. Prediction overlap was assessed by correlating the actual at-risk t-statistic vector with the predicted t-statistic vector, after confining to the most highly dysfunctional voxels in the actual data (identified by thresholding the actual t-statistic vector at p<.001 uncorrected). Significance was assessed by a non-parametric permutation approach, wherein a null overlap distribution was generated by randomly selecting a matched number of voxels from the predicted t-statistic vector and correlating with the (unscrambled) actual thresholded t-statistic vector over 1000 iterations. Significance of the observed overlap r values was assessed against this null distribution at an alpha level of .05.

### Statistical analyses

Statistical analyses conducted for this report have been described in detail in preceding Method sections and in the Results section. To summarize, significance of the classification accuracies in the machine learning analyses was estimated parametrically via binomial test against chance 50% accuracy. Significance of the overlap between the activity flow-predicted task activations and the actual activations was assessed at both the group and subject level, via both parametric (group-level: p value accompanying the overlap r; subject-level:one-sample t-test for Fisher-transformed overlap r values against 0) and non-parametric permutation test approaches (see Method for details). Similar group/subject and parametric/non-parametric approaches were adopted for the activity flow analysis predicting task activation dysfunction via regression. Significance of the overlap between the activity flow-predicted contrast images and the actual contrast images was assessed via a non-parametric permutation test controlling for inflation of the overlap r value by the subtraction operation (see previous section).

We also conducted a series of follow-up analyses probing the extent to which activity flow mapping had transformed healthy task activations into unhealthy at-risk ones (central to the theoretical basis of the framework, see Figure 1a), as well as the extent to which the predicted activations captured task-specific and subject-specific features of the actual activations. These analyses are more fully described in the relevant Results sections, but operate on a similar analytic principle of contrasting the similarity (Pearson correlation) between at-risk AD subjects’ predicted task activations and appropriate activation templates via paired-sample t-tests (e.g. for the transformation analyses: contrasting the similarity of at-risk subjects’ predicted activations to healthy versus actual at-risk group templates). We also conducted a series of control analyses which are fully described in the last section of the Results.

## Results

### Evidence of at-risk Alzheimer’s-related dysfunction: behavioral

Consistent with their at-risk AD status, none of the included subjects met clinical criteria for full Alzheimer’s-related behavioral impairment: Mini Mental State Exam (MMSE) scores were greater than 24, and Clinical Dementia Ratings (CDR) were 0 (negative dementia diagnosis). However, analysis of the MMSE scores revealed that the at-risk group was behaviorally impaired compared to the healthy group, even in this non-clinical range (see Table 1 for MMSE means and standard deviations across groups). MMSE scores were significantly lower for the at-risk group across both APOE (2-sample t-test healthy > at-risk: t(99)=2.32, p=.02) and amyloid (t(99)=2.20, p=.03) segregation types. This supports the presence of meaningful Alzheimer’s-related dysfunction in the at-risk AD subjects, which activity flow mapping attempted to capture.

### Evidence of at-risk Alzheimer’s-related dysfunction: restFC and task activation classifications

To further demonstrate dysfunction in the at-risk AD versus healthy group, we trained multivariate machine learning classifiers to discriminate between the two groups separately on the basis of regional restFC and regional task activation features. We used an RSA-based classifier with leave-one-subject-out cross-validation (see Method). To increase the sample size for classifier training, these analyses collapsed across at-risk subjects identified in either APOE and amyloid segregation types (yielding at-risk n=40).

For the restFC classification, we confined features to the connections in the full restFC matrix that most strongly discriminated between the two groups, as identified in the training set (see Method for details; Dosenbach et al., 2010; Vergun et al., 2013). The restFC classifier achieved significantly above-chance accuracy in discriminating between at-risk and healthy groups (66.3% overall accuracy; 70% sensitivity in classifying ‘at-risk’, 62.5% specificity in classifying ‘healthy’; p=.002 via binomial test). Classifiers trained to discriminate using task activation features yielded similar accuracy, achieving 68.8% accuracy for the animacy task (75% sensitivity, 62.5% specificity, p=.001) and 58.8% for the Stroop task (62.5% sensitivity, 55% specificity, p=.073). A separate classifier was trained to discriminate on the basis of ‘task general’ activation features identified by averaging across the two tasks for each subject, yielding significantly above-chance accuracy (65% overall accuracy; 67.5% sensitivity, 62.5% specificity, p=.005). Note that performance of the restFC and task activation classifications was similar when using a leave-two-subject-out cross-validation approach, with one unhealthy and one healthy subject held-out to ensure perfectly balanced group sizes on each training loop: restFC classification accuracy=60%, p=.047; animacy task activation accuracy=66.3%, p=.002; Stroop task activation accuracy=56.3%, p=.157; task general activation accuracy=66.3%, p=.002).

Overall, the classification results highlight reliable differences between the at-risk and healthy groups in terms of their restFC and task activations. Considered with the between-group MMSE differences reported above, this validates the presence of meaningful dysfunction in the at-risk AD group, and hence supports our theoretical approach of applying activity flow mapping to model this dysfunction.

### Activity flow mapping accurately predicts task activation patterns in held-out at-risk AD subjects

Activity flow mapping was firstly applied to predict at-risk AD task activations at the region level (Figure 3a). For the APOE segregation, the group-level predicted-to-actual activation overlap was significant for both the animacy task (Pearson r=.68, p<.00001) and the Stroop task (r=.70, p<.00001). Significance was maintained when computing the overlap at the subject level (treating subjects as a random effect, see Methods) for both the animacy task (mean r=0.24, t(32)=7.80, p<.00001) and the Stroop task (mean r=0.18, t(32)=6.84, p<.00001). Visual inspection of the commonalities across the predicted and actual regional vectors (see Figure 3a) highlights the cross-task recovery of the canonical task-negative deactivation of the default mode network (DMN) and task-positive activation of the cognitive control networks (CCNs; including the frontoparietal control network, FPN, cingulo-opercular network, CON, dorsal and ventral attention networks, DAN and VAN).

**Figure 3.**
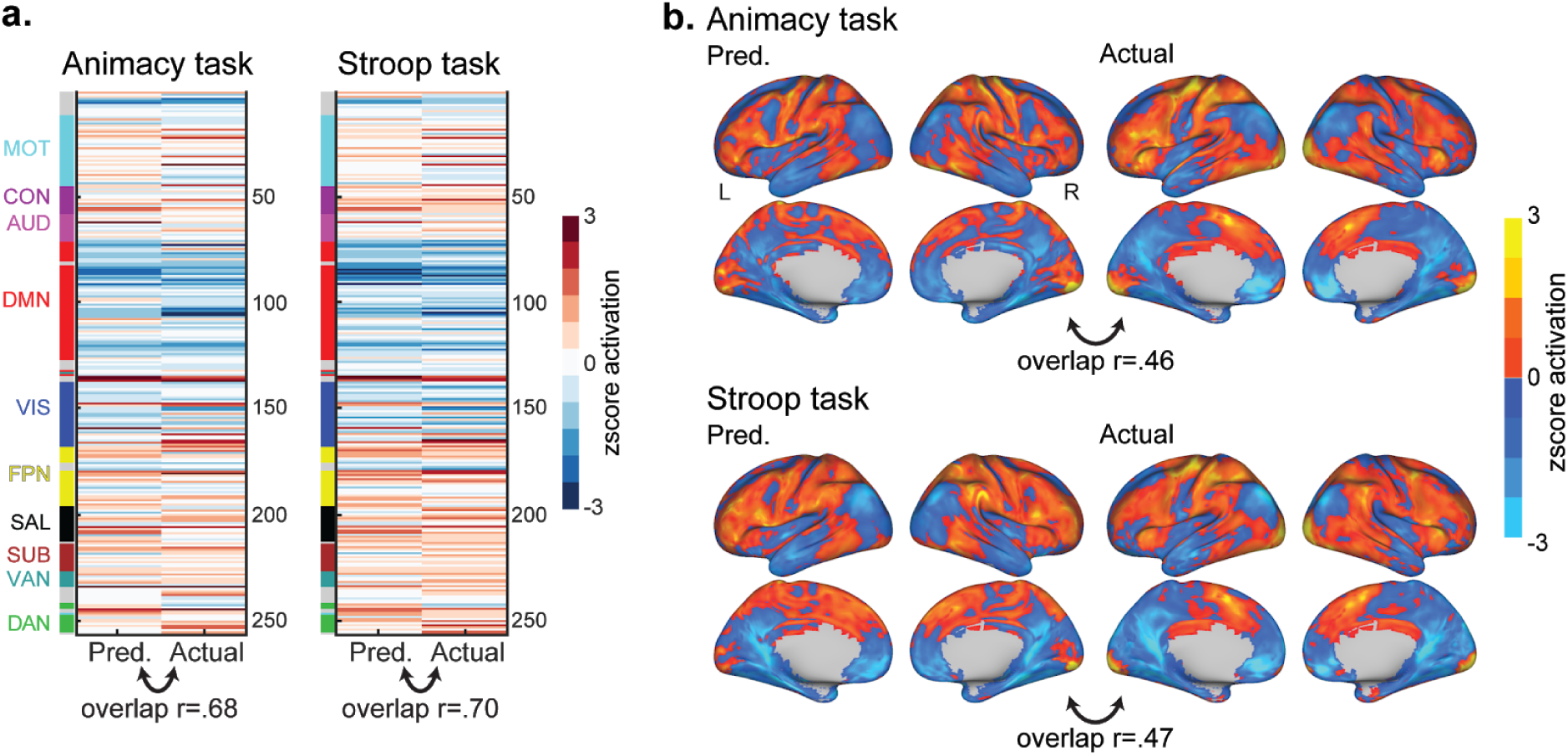
Activity flow mapping results for the prediction of task activations in at-risk AD subjects, as identified by the APOE segregation. **a)** Group-averaged predicted and actual task activation vectors for the regionwise analysis, and their overlap (Pearson r). Network affiliations for each region from the Power et al (2011) atlas are provided in the color-coded bar to the left: MOT=motor network, CON=cingulo-opercular network, AUD=auditory network, DMN=default mode network, VIS=visual network, FPN=fronto-parietal control network, SAL=salience network, SUB=subcortical network, VAN=ventral attention network, DAN=dorsal attention network. **b)** Group-averaged predicted and actual task activation brain maps for the voxelwise analysis, and their overlap (Pearson r). All regionwise and voxelwise activations were zscored for visualization purposes.

Non-parametric permutation tests were also conducted to demonstrate the importance of region-to-region correspondence between the activation and restFC terms to the activity flow predictions, and deal with any violation of the independence assumption in the parametric tests (see Methods). The pattern of results obtained with this non-parametric approach was identical to that obtained with parametric methods: for the APOE segregation, prediction overlap was significant at both the group level (p<.001 for both tasks) and the subject level (p<.001 for both tasks).

Activity flow mapping was extended to predict task activations at every voxel in the brain (Figure 3b). To prevent circularity arising from spatial autocorrelation of fMRI data, voxels within the same functional region as the to-be-predicted region were excluded as well as all voxels within a 9mm radius of that region (see Method). The overlap between the group-averaged predicted and actual voxelwise activations was significant for both tasks: animacy r=.46, p<.00001; Stroop r=.47, p<.00001. Significant prediction accuracy was maintained when computing the voxelwise overlap at the subject level (as per the regionwise analyses above): animacy r=.13, t(32)=6.20, p<.00001; Stroop r=.09, t(32)=4.76, p<.0001. We also conducted permutation tests for the voxelwise analyses following the same rationale as the regionwise results. This yielded an identical pattern as the parametric voxelwise results: prediction overlap was significant at both the group level (p<.001 for both tasks) and the subject level (p<.001 for both tasks). Visual inspection of the common activation patterns across the predicted and actual voxelwise brainmaps (see Figure 3b) again demonstrates cross-task recovery of canonical task negative (DMN regions e.g. bilateral precuneus, posterior cingulate, angular gyrus and ventromedial prefrontal cortex) and task positive (CCN regions e.g. bilateral anterior cingulate, superior frontal gyrus, lateral prefrontal cortex and inferior parietal cortex) activations by the activity flow mapping procedure.

Similarly high prediction accuracy was obtained when activity flow was applied to at-risk AD subjects identified by the amyloid segregation (see final Results section). Overall, the results demonstrate accurate prediction of task activations in held-out at-risk subjects by activity flow mapping, as estimated at the group and subject level, at regionwise and voxelwise spatial scales, across two distinct cognitive tasks and across two distinct ways of identifying at-risk AD subjects.

### Activity flow over at-risk AD restFC transforms a healthy aged activation state into an unhealthy one

We next sought more direct evidence that activity flow over an at-risk pattern of restFC transforms a healthy activation state into an at-risk one (see Figure 1a). To clarify, the extent to which the activity flow model makes accurate predictions of at-risk AD activations is determined by how successful the model’s alteration of the healthy activation template by at-risk restFC is in transforming that template into the actual at-risk activation. We formally tested this transformation hypothesis by comparing the similarity between each at-risk subject’s predicted activation with the group healthy activation versus the actual group at-risk activation (see Figure 4a). If the similarity of the predicted-to-actual at-risk activation was reliably greater than the similarity of the healthy-to-actual at-risk activation, this would support a mechanistic link between altered restFC and the emergence of dysfunctional task activations. The actual group at-risk activation was averaged after excluding the to-be-compared at-risk subject. This yielded a pair of Pearson r values for each subject (at-risk task similarity r and healthy task similarity r, averaged across the two tasks), which were Fisher-transformed and compared via a paired-sample t-test.

**Figure 4.**
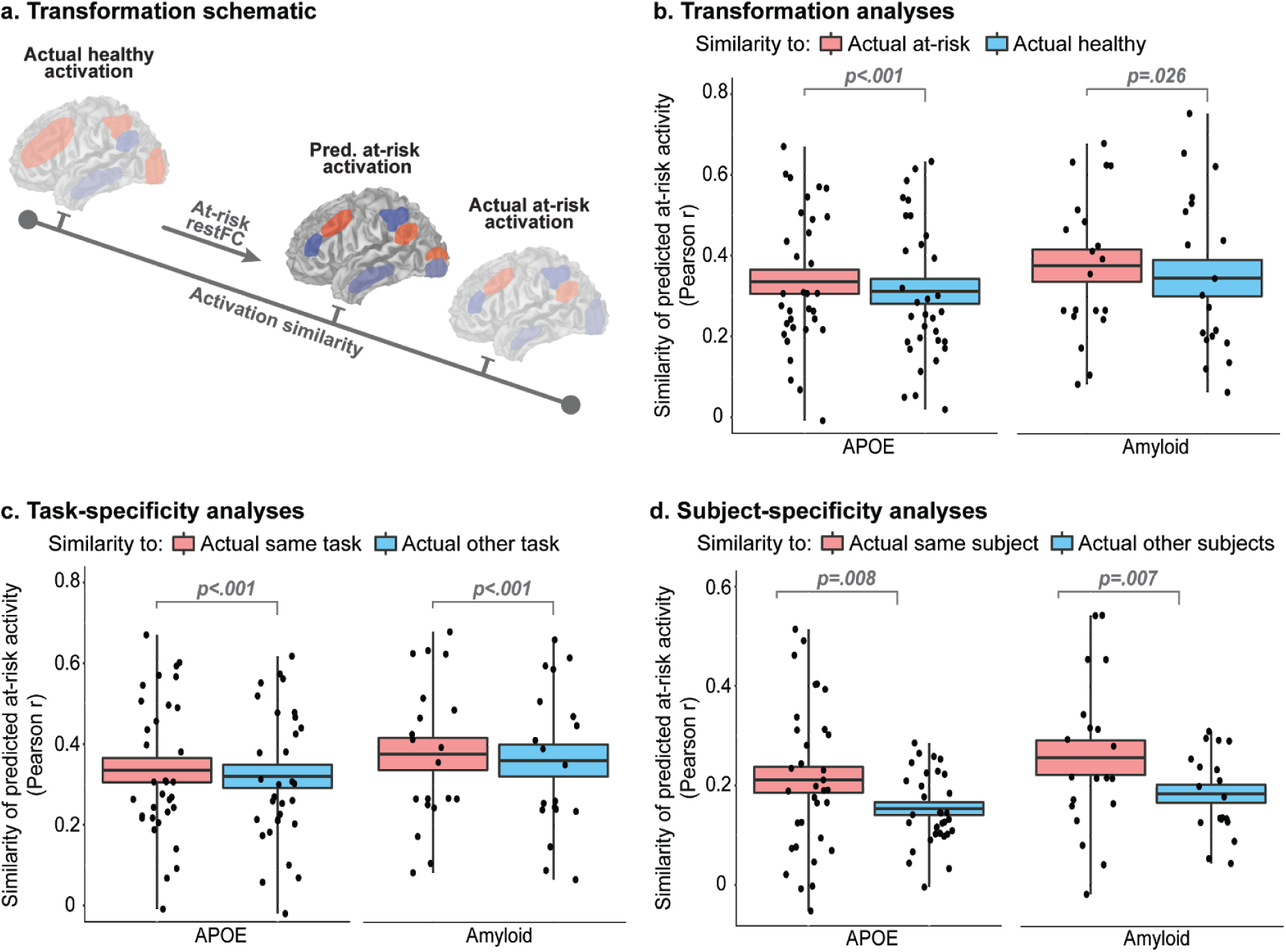
Activity flow mapping recovers salient task activation properties in at-risk AD subjects. **a)** Schematic demonstration of the transformation analysis: operation with at-risk subject restFC transforms a healthy task activation into an at-risk unhealthy one, formalized as greater similarity with the actual at-risk group activation template versus the healthy group template. Panels b-d follow a similar approach of comparing the similarity (Pearson r) between the predicted at-risk subject activations with two activation templates (denoted for each analysis by the legend in each panel), followed by group-level contrasts of the resulting two similarity values (after Fisher-z transform) via paired t-test. All plots represent the mean +/− the standard error of the similarity r values (the box) and the minimum/maximum (the whisker), with individual data points overlaid (r values for each at-risk subject). All panels depict results across the APOE (left plot) and amyloid (right plot) segregation types. **b)** Results for the transformation analyses: the similarity between the activity flow-predicted at-risk activations for each subject was compared with group-averaged templates of the actual at-risk and actual healthy activations. **c)** Activity flow mapping captures activation patterns that are specific to a given task. Similarity of predicted at-risk activation to actual at-risk activation from the same task was compared to similarity with the actual at-risk activation from the other task (both group templates). **d)** Activity flow mapping captures activation patterns that are specific to a given subject. The similarity of the predicted activation for a given at-risk subject to the actual activation for that subject was contrasted with similarity to the actual activations for the other at-risk subjects (averaged across subjects).

As expected, the predicted at-risk activations were significantly more similar to the actual at-risk group activation than to the actual healthy group activation (Figure 4b). This held for both the APOE segregation (actual at-risk > actual healthy similarity, paired-sample t(32)=4.74, p<.001) and the amyloid segregation (t(19)=2.41, p=.026). These results are noteworthy given that the group healthy activation vector is the only activation term the model has access to when predicting at-risk AD subject activations (see Figure 1b). That the activity flow-predicted activations are actually more similar to the actual at-risk group activation hence provides compelling evidence of the power of restFC alone in transforming healthy into unhealthy activation states.

### Activity flow mapping activation predictions are task-specific

To further validate the accuracy of activity flow mapping, we interrogated whether the predicted activations captured features specific to the to-be-predicted task. This property is particularly desirable when applied in clinical contexts, as it enables prediction of brain activations in different tasks designed to isolate different cognitive processes and associated impairment (rather than solely capturing task-general activation patterns).

For each at-risk AD subject, we compared the similarity (Pearson r) of their predicted activations to the group-averaged actual at-risk activation for the same task with the similarity to the group-averaged actual at-risk activation for the other task. To clarify, to conduct the analysis for the animacy task, we would compute the similarity of the predicted animacy activation for an at-risk subject to the group-averaged actual at-risk animacy activation (same task), and compare this to the similarity to the group-averaged actual at-risk Stroop activation (other task). Both at-risk group activation templates (same task and other task) excluded the at-risk subject from which the predicted activation is taken. This yielded a pair of Pearson r values for each subject (same task similarity r and other task similarity r, each averaged across the two tasks), which were Fisher-transformed and contrasted via paired-sample t-test.

The results reveal that the activity flow-predicted at-risk subject activations were significantly more similar to the actual at-risk group activation for the same task than the other task (see Figure 4c for mean similarity values). This held for both the APOE segregation (same > other task similarity, paired-sample t(32)=5.80, p<.001) and the amyloid segregation (t(19)=5.89, p<.001).

### Activity flow mapping activation predictions are subject-specific

As another demonstration of prediction accuracy and potential clinical utility, we sought evidence that the activity flow-predicted activation for a given at-risk subject was more similar to the actual activation for that same subject versus the activation of other at-risk subjects in the sample. For each at-risk AD subject we compared the similarity between their predicted task activation and their actual task activation (same subject similarity) versus the similarity with the actual task activations of all other at-risk subjects (other subject similarity). The similarity r values computed with the other subjects were averaged to yield one ‘other subject’ similarity r value, which was contrasted with the ‘same subject’ similarity r value via paired t-test (after Fisher-z transform).

The results confirmed that the activity flow-predicted task activation for a given at-risk subject was significantly more similar to their own actual activation pattern (see Figure 4d for mean similarity values): APOE segregation (same > other subject similarity, paired t(32)=2.85, p=.008), amyloid segregation (t(19)=3.04, p=.007). This illustrates the ability of the activity flow framework to capture individualized aspects of at-risk AD task activation patterns. This again enhances the framework’s potential clinical utility, given considerable recent biomedical research interest in modeling individual differences to develop personalized medicine interventions (Matthews and Hampshire, 2016).

### A second cycle of transformation via restFC improves activity flow mapping predictions

As an extension to the idea that restFC transforms healthy activation states into unhealthy ones (see earlier Results section), we interrogated whether one cycle of transformation as applied thus far is optimal or whether a second cycle would improve prediction accuracy. This idea is inspired by the similarity of activity flow mapping to simulation of a single propagation ‘time step’ in a neural network model (McClelland and Rogers, 2003; Cole et al., 2016), suggesting simulation of an additional time step might (despite fMRI-estimated dynamics being relatively slow) result in additional transformation of activation patterns. Specifically, after generating a predicted task activation for an at-risk AD subject, we passed that predicted task activation through the model a second time i.e. multiplying the predicted task activation vector with the restFC vector for that subject (see Figure 5a).

**Figure 5.**
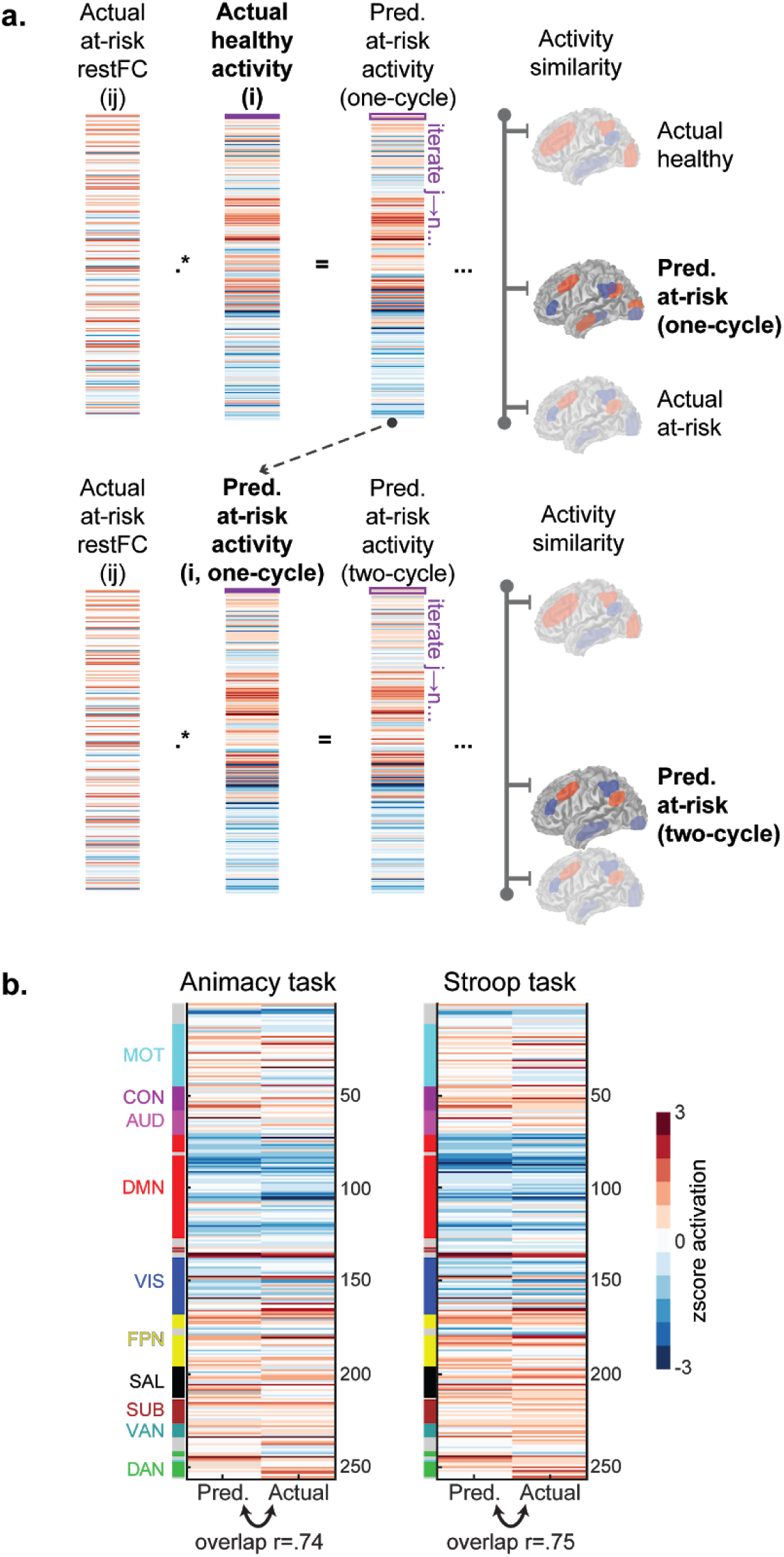
Demonstration and results for two-cycle activity flow mapping. **a)** Conceptual schematic depicting the two-cycle approach. In the standard one-cycle approach (upper panel, also illustrated in Figure 1b and figure 2c), at-risk AD activations are generated for a to-be-predicted region (j) from the dot product of the subject’s restFC to region j (ij) and the whole-brain activation state (i) estimated from the healthy group mean. Iterating through all values of j for all regions/voxels (n) populates the predicted at-risk activation vector (upper panel, third vector). This first cycle of activity flow mapping transforms the healthy activation into an at-risk activation to some degree, by increasing the similarity of the predicted with the actual at-risk activation (upper right). In the two-cycle step that follows (lower panel), the one-cycle-predicted at-risk activation is again multiplied with the subject’s rest FC, supplanting the group healthy activation that generated the one-cycle prediction. For these analyses, the overlap with the actual at-risk subject/group activations is hence computed with the two-cycle-predicted activation, which theoretically should improve the similarity with the actual at-risk activations (lower right). **b)** Results of the two-cycle activity flow mapping approach support this hypothesized improvement. The one-cycle prediction overlap results for comparison: r=.68 for animacy and r=.70 for Stroop. Plotting conventions are the same as in Figure 3a.

As expected, this two-cycle transformation approach improved the overlap between regional predicted and actual activations. For the APOE segregation, the group predicted-to-actual activation overlap was significant for both the animacy task (pearson r=.74, p<.00001; r increase versus standard one-cycle activity flow=.06) and the Stroop task (r=.75, p<.00001; r increase versus one-cycle=.05). Significant overlap was maintained when treating the subjects as a random effect, for both the animacy task (mean r=0.27, p<.00001; r increase versus one-cycle=.03) and the Stroop task (mean r=0.22, p<.00001; r increase versus one-cycle=.04). This improvement in the subject-level overlap was found to be significant via paired t-tests for both the animacy task (two-cycle > one-cycle, t(32)=3.47, p=.002) and the Stroop task (t(32)=5.53, p<.001). The two-cycle model predictions also captured similar properties of restFC-induced transformation, task-specificity and subject-specificity to the original model, as described in the preceding sections.

The two-cycle model also improved prediction accuracy in the voxelwise analyses. The overlap between the group-averaged voxelwise predicted and actual task activations was significant for both tasks: animacy r=.56, p<.00001 (r increase versus one-cycle=.10); Stroop r = .61, p<.00001 (r increase versus one-cycle=.14). Significant prediction overlap was maintained at the subject level: animacy r=.16, p<.00001 (r increase versus one-cycle=.03); Stroop r=.14, p<.0001 (r increase versus one-cycle=.05). The improvement in subject overlap was found to be significant via paired t-tests for both the animacy task (two-cycle > one-cycle, t(32)=7.25, p<.001) and the Stroop task (t(32)=8.40, p<.001).

Overall, results from the two-cycle activity flow model further support the mechanistic underpinnings of the approach, given the closer approximation of the two-cycle models (despite fMRI-estimated dynamics being relatively slow) to fully dynamic neural simulations with many cycles/time steps. Further, the subject level prediction accuracy improvement suggests clinical utility by better capturing individualized activation features.

### Isolation of dysfunctional task activation predictions by regressing out healthy activations

Thus far, we have demonstrated the high accuracy of activity flow mapping when used to predict task activations in at-risk AD subjects. This prediction did not directly identify how the regionwise/voxelwise activation patterns differed between at-risk and healthy subject groups. Demonstrating that activity flow mapping is able to accurately capture such differences is desirable for the aim of modeling dysfunction in AD (and other pathologies). To achieve this, we regressed out the group healthy task activation from both the predicted and actual task activations for each at-risk subject (see Method for details). The predicted-to-actual activation overlap computed on the residual vectors hence indexed the ability for activity flow to capture differences between the healthy group and that at-risk subject.

For the APOE segregation, the overlap between predicted and actual dysfunctional activations was significant for both the animacy task (Pearson r=.35, p<.00001) and the Stroop task (r=.32, p<.00001). Significant overlap was maintained when treating the subjects as a random effect, for both the animacy task (mean r=.13, p<.00001) and the Stroop task (mean r=.07, p=.004). As with the earlier activity flow mapping results, we also assessed the significance of the activity flow ‘dysfunction’ predictions via non-parametric permutation tests (see Method for details). This yielded an identical pattern of results to the parametric statistics: dysfunctional prediction overlap was significant at the group level (p<.001 for both tasks) and the subject level (p<.001 for the animacy task, p=.005 for the Stroop task). The pattern of results was identical for the amyloid segregation type (see final Results section).

We also predicted task activation dysfunction at every gray matter voxel in the brain. The voxelwise overlap between predicted and actual dysfunctional activations was significant for both the animacy task (Pearson r=.29, p<.00001) and the Stroop task (r=.18, p<.00001). Significant overlap was maintained when treating the subjects as a random effect for both the animacy task (mean r=.09, p<.00001) and the Stroop task (mean r=.07, p<.001). The pattern of significance was identical via non-parametric permutation tests (following the same rationale as the regionwise analyses), for both the group-level (p<.001 for both tasks) and subject-level overlap (p<.001 for both tasks). Overall, these findings support the ability of activity flow mapping to capture task activation differences between the at-risk and healthy groups, suggesting utility in modeling AD-related dysfunction.

### Prediction of between-group contrast images also captures task activation dysfunction

The preceding section described our use of activity flow mapping to predict at-risk AD task activations directly linked to clinical dysfunction. As an alternative to the regressing-out approach, we generated between-group contrast images capturing voxelwise task activation differences between the at-risk and healthy subjects (see Method). This was done both using the activity flow-predicted at-risk activations (‘predicted contrast image’=group average predicted at-risk minus group-average healthy) and the actual at-risk activations (‘actual contrast image’=group average actual at-risk minus group-average healthy). The predicted-to-actual contrast image overlap was quantified via Pearson correlation as before. A non-parametric permutation approach was used to assess significance whilst controlling for any inflation of the overlap correlation by the subtraction of the group healthy activation (see Method).

For the APOE segregation, the overlap between the predicted and actual contrast images was r=.39 and r=.30 for the animacy and Stroop tasks respectively. Both overlap correlations were significant versus the permuted null (p<.001 for both tasks). A thresholded version of these analyses confined the predicted-to-actual contrast image overlap to only those voxels that significantly differed in the actual contrast image map (via 2-sample t-test at p<.001 uncorrected, see Method for details). This also yielded significant overlap: animacy r=.53, p<.001 (via non-parametric permutation test, see Method); Stroop r=.67, p<.001. This demonstrates that activity flow mapping is able to accurately predict the most highly dysfunctional voxelwise task activations in the at-risk AD subjects. Overall, the generation of between-group contrast images via activity flow mapping raises another potentially fruitful clinical application by providing visually interpretable dysfunction brainmaps.

### Generalization and robustness of the at-risk AD activation predictions

Firstly, all results held when segregating subjects into at-risk and healthy groups on the basis of high levels of PET amyloid beta rather than presence of APOE *ε*4. Significant group-level overlap was obtained for the main regional activity flow analyses (as described in Results section ‘Activity flow mapping accurately predicts task activation patterns in held-out at-risk AD subjects’): animacy group r=.70 and Stroop group r=.68 (both p<.00001; see Figure 7a). Significant overlap was also obtained when confining the regional predictions to dysfunctional activations (as described in Results section ‘Isolation of dysfunctional task activation predictions by regressing out healthy activations’): animacy group r=.40 (p<.00001) and Stroop group r=.25 (p<.0001). Prediction accuracy held when overlap was computed at the subject level (not presented), and also for all major voxelwise analyses (see Figure 7b for the main voxelwise results). Prediction accuracy also held when identifying ‘purer’ segregations of at-risk subjects, by excluding subjects that had both high levels of PET amyloid and presence of APOE *ε*4 from either segregation. Activity flow mapping yielded significant prediction accuracy in these purer segregations despite the reduction in sample size (at-risk n for pure APOE=20; n for amyloid=7), for both the pure APOE segregation (animacy group r=.62, Stroop r=.65, both p<.00001) and the pure amyloid segregation (animacy group r=.58, Stroop r=.58, both p<.00001). Note also that concatenating at-risk subjects that were identified either in amyloid or APOE segregation types also yielded high prediction accuracy in the main analyses (combined at-risk n=40): animacy group overlap r=.71 and Stroop r =.74 (both p<.00001). Collectively, these findings highlight the generalizability of activity flow mapping in predicting AD-related dysfunction across different ways of identifying at-risk subjects.

**Figure 6.**
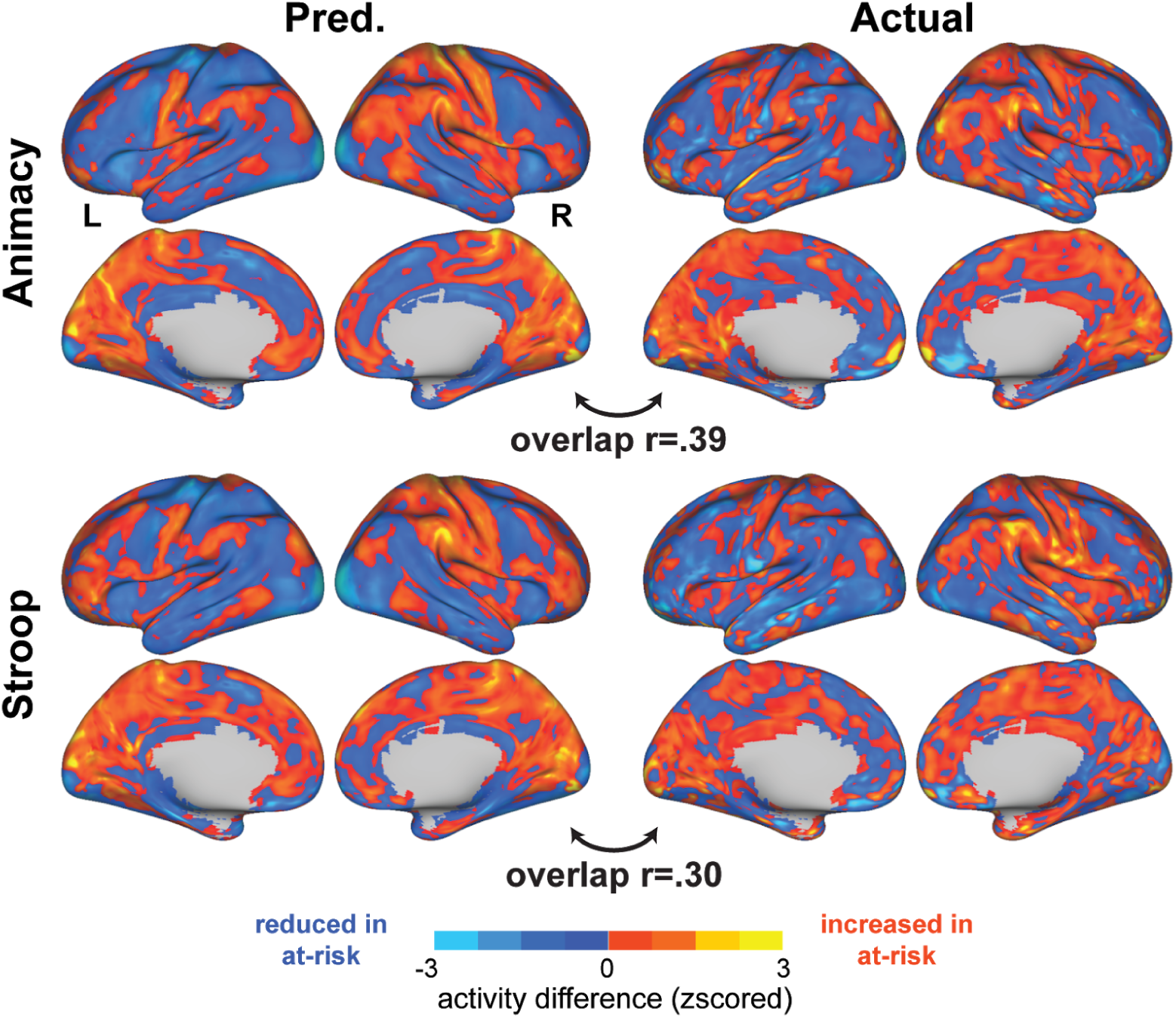
Generation of between-group activation contrast images via activity flow mapping. Predicted (left panel) and actual (right panel) contrast images for the animacy task (upper panel) and Stroop task (lower panel) are depicted for the APOE segregation, along with the predicted-to-actual contrast image overlap (both r values p<.001 via permutation test, see Results for details). Activation differences were generated as group averaged predicted/actual at-risk AD minus group averaged healthy, and were zscored for visualization purposes.

**Figure 7.**
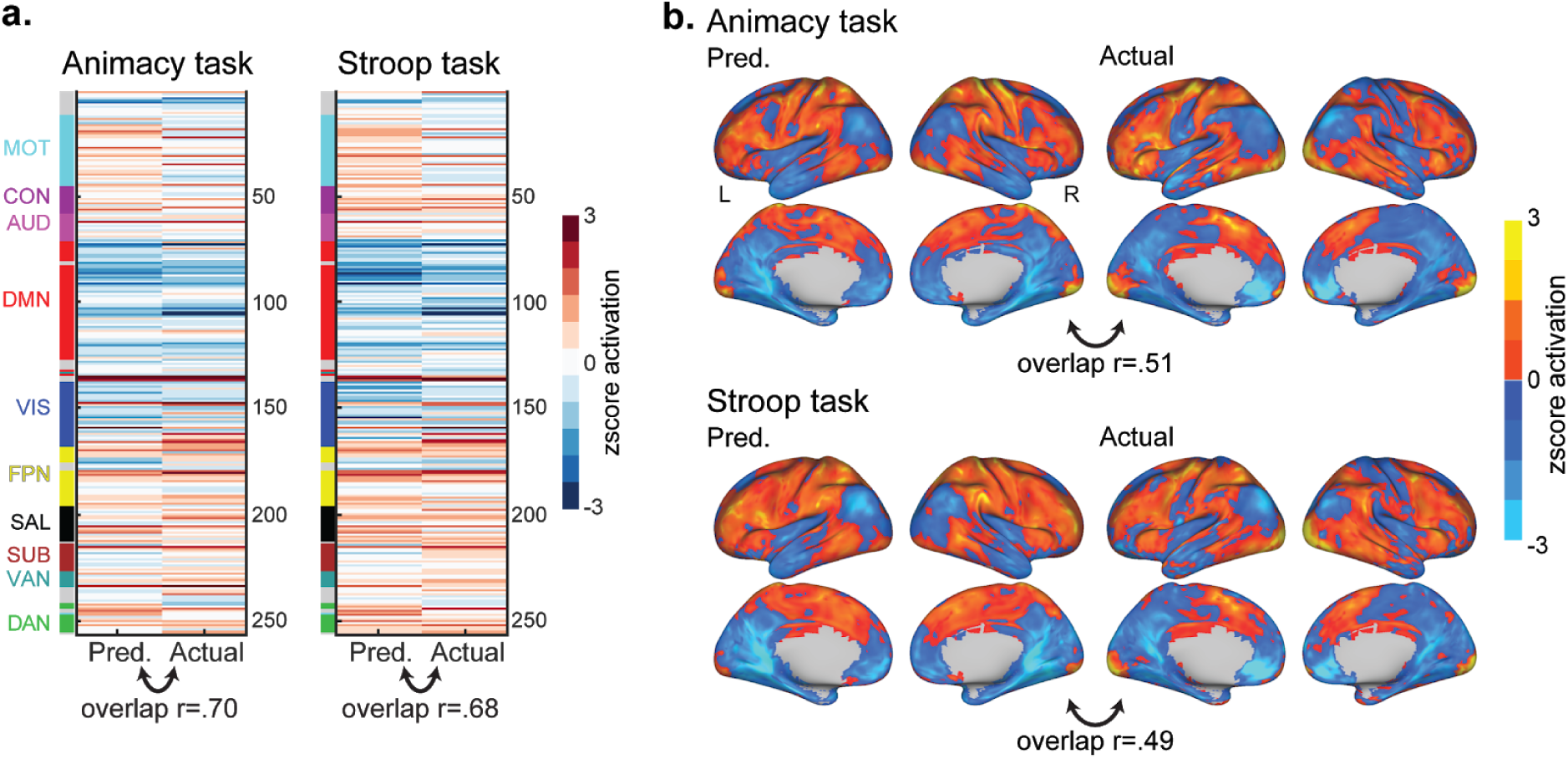
Activity flow mapping results for the prediction of task activations in at-risk AD subjects, as identified by the amyloid segregation. **a)** Predicted and actual task activation vectors for the regionwise analysis and **b)** the voxelwise analysis, displayed with the group-level predicted-to-actual overlap. Plotting and statistical conventions are identical to Figure 3.

The high prediction accuracy of activity flow mapping was maintained when performing global signal regression prior to computing restFC: animacy group overlap r=.70 and Stroop r=.74 (both p<.00001) for the APOE segregation. The results also held when estimating restFC via Pearson correlation rather than multiple regression FC: animacy group overlap r=.57 and Stroop r=.63 (both p<.00001) for the APOE segregation. This shows robustness of the approach to variation in restFC preprocessing and estimation.

We also probed potential influences of vascular (i.e. non-neural) components of the fMRI BOLD signal on the accuracy of the activity flow mapping predictions. To estimate vascular BOLD contributions at each region, we used the resting-state fluctuation amplitude (RSFA), computed as the temporal standard deviation across resting-state timepoints, given previously established links between RSFA and vasculature (Kannurpatti and Biswal, 2008; Tsvetanov et al., 2015; Mayhew et al., 2016). The regional RSFA estimates for each at-risk subject were regressed out of their actual task activations and the activity flow-predicted activations, prior to computing subject-level overlap (Pearson’s r). Overlap between predicted and actual activations remained significant after this operation, for both the APOE (animacy r=.21, Stroop r=.17; both p<.00001) and amyloid (animacy r=.23, Stroop r=.22; both p<.0001) segregations. This addresses the possibility that the accuracy of activity flow mapping is driven by vascular rather than neural components of the BOLD signal, which is important given prior demonstration of vasculature confounds of age-related task activation effects (Kannurpatti et al., 2011; Tsvetanov et al., 2015)

Finally, we demonstrated that age, gender and motion did not appreciably influence the accuracy of activity flow predictions. The correlation between age of the at-risk AD subjects and their subject-level predicted-to-actual activation overlap was non-significant for both tasks, both for the APOE (animacy r=-.23, p=.21; Stroop r=-.18, p=.31) and amyloid (animacy r=-.09, p=.71; Stroop r=-.13, p=.58) segregations. We probed the effect of gender by conducting a 2-sample t-test contrasting the subject-level predicted overlap r values (after fisher transformation) between female and male subjects. This verified that the strength of activity flow mapping was not significantly influenced by gender, both for the APOE (animacy female > male t(31)=0.02, p=.98; Stroop t(31)=0.75, p=.46) and amyloid (animacy t(18)=-0.72, p=.48; Stroop t(18)=0.76, p=.46) segregations. Finally, to probe any effect of subject motion, we correlated the mean and peak framewise displacement for each subject (FD; Power et al., 2012), estimated separately for rest and task sessions, with their subject-level predicted activation overlap. Across mean/peak measures of FD, estimated across rest/task runs and across amyloid/APOE segregations, no significant relationship between in-scanner motion and the accuracy of activity flow predictions was found (p accompanying the r values >.1 in all cases).

## Discussion

We report the first extension of the activity flow mapping framework (Cole et al., 2016) to a clinical context: modeling dysfunction in at-risk AD subjects. The framework was applied in an elderly sample segregated into at-risk and healthy subgroups based on established AD risk factors (presence of APOE *ε*4 or high PET amyloid). Activity flow mapping predicted activations in held-out at-risk subjects (and held-out brain regions/voxels) from their individual pattern of restFC altering a healthy task activation template. This procedure accurately predicted at-risk AD activations across different cognitive tasks, neural spatial scales, and at-risk segregation types, illustrating its robustness. Follow-up analyses confirmed key mechanistic properties of our framework: healthy activations were transformed by the individual pattern of at-risk restFC alone, and this transformation was improved by a second cycle. In support of the framework’s clinical utility, activation predictions were statistically reliable at the individual level, and captured task-specific and subject-specific signatures. Finally, activity flow mapping remained accurate when predicting overtly dysfunctional activations (those that differed for at-risk versus healthy subjects).

Our findings have three broad implications. Firstly, the success of activity flow mapping provides empirical evidence for a mechanistic role of restFC in defining the pathways over which activity flows across the brain and, therefore, pathways underlying activity dysfunction. Secondly, our findings highlight the potential of applying activity flow mapping to predict AD-related pathology. Thirdly, these results provide theoretical insights that can guide ongoing efforts to optimize the use of restFC to predict/classify disease features more generally.

The first implication relates to much needed insight into the cognitive relevance of restFC, which has been questioned given the experimentally unconstrained nature of rest (Mill et al., 2017). A role for connectivity in driving the emergence of task activations has long been formalized in computational models (Hodgkin and Huxley, 1952), and empirically demonstrated by local circuit neurophysiology linking connectivity to synaptic modulation of action potentials (Laughlin and Sejnowski, 2003; Fries, 2005). A similar role for synaptic connectivity in weighting paths of information propagation across computational ‘layers’ has been a unifying property of neural network simulations of cognitive function (McClelland and Rogers, 2003; Battaglia et al., 2012). Importantly, the success of activity flow mapping suggests restFC as a large-scale analogue of such synaptic processes, corroborating its utility in mapping intrinsic network organization (Raichle, 2010). By extension, incorporating network connectivity is likely central to achieving a full theoretical understanding of how the brain activates during cognitive processing. This goes against the historical isolation of connectivity and task activation studies into separate neuroscientific subfields, and instead promotes their mechanistic unification via activity flow. Future research applying activity flow mapping to more direct measures of neural activity than the fMRI BOLD response (e.g. human electrocorticography or source-modeled electro-/magnetoencephalography) could substantiate this association between restFC and synaptic processes.

Our findings build on previous successful demonstrations of activity flow mapping (Cole et al., 2016; Ito et al., 2017) by extending the approach to predicting activations and activation-related dysfunction in a clinical context. Our focus on AD was motivated by its grim prognosis, raising an urgent need to develop imaging biomarkers that can aid early diagnosis and intervention. It was also motivated by the strong evidence base (relative to other disorders; Fox and Grecius, 2010; Woo et al., 2017) characterizing restFC and task activation alterations in at-risk (Hedden et al., 2009; Sheline and Raichle, 2013; Oh et al., 2016) and later stages of AD (Grady, 2012; Ferreira and Busatto, 2013; Campbell and Schacter, 2017). Considered with the accuracy of our restFC-based machine learning classification of at-risk subjects, the success of activity flow in the present at-risk sample strengthens previous staging models of AD risk factors (Jack et al., 2010; Sheline and Raichle, 2013). These suggest that restFC alterations emerge many years prior to clinical impairment. Critically, rather than studying restFC alterations separately from task activations, the activity flow framework unifies both measures mechanistically (Figure 1). This affords a synthesis of the practical benefits of acquiring resting-state data (e.g. reducing overall scanner time, enabling imaging assays for clinical samples unable to perform in-scanner tasks) with clearer theoretical grounding via linking to task activations and associated cognitive function.

Future research will need to clarify the chronology of restFC alterations relative to the emergence of PET amyloid and other AD signatures (e.g. tau deposition; Jones et al., 2016, 2017). Relatedly, recent demonstration that connectivity at an earlier timepoint (5 years of age) predicts the location of fMRI task activations associated with later development of reading ability (assessed at 8 years of age; Saygin et al., 2016), suggests potential to use activity flow to predict task activation dysfunction well in advance of its actual emergence. This would have clear clinical utility, and calls for the extension of the activity flow framework to predict timecourse features of AD (i.e. onset, prognosis). A prerequisite to this endeavour is the furthering of recent attempts (e.g., the Alzheimer’s Disease Neuroimaging Initiative) to collect longitudinal data in large elderly samples spanning the full spectrum of AD-related pathology: healthy aged, at-risk, MCI, and post-AD onset. A related future refinement to the activity flow model will be to link its predictions to behavior and, importantly for clinical applications, behavioral impairment.

In interrogating the general utility of activity flow for clinical neuroscience, another promising avenue would be to extend the approach to pathologies beyond AD. Theoretically, activity flow modeling can be applied to any meaningful categorization of unhealthy and healthy subgroups, given that it makes predictions on the basis of the unhealthy subjects’ restFC altering a healthy group activation template. Notably, the mechanistic basis of the activity flow framework (Cole et al., 2016) differentiates it from other more data-driven ‘fingerprinting’ approaches with similar predictive aims (Rosenberg et al., 2016; Saygin et al., 2016; Tavor et al., 2016; Lin et al., 2018). These approaches use abstract coefficients to translate an individual’s connectivity profile into predicted task activations (or behavior), as estimated via intensive optimization/cross-validation during model training. Such abstract data-driven provisions convey a lack of theoretical grounding (‘neuroscientific validity’), which has been highlighted as an obstacle to integrating connectivity-based predictive models into clinical practice (Stephan et al., 2015; Matthews and Hampshire, 2016; Woo et al., 2017). In contrast, the activity flow model assumes that the whole-brain activation state translates a connectivity profile into a task activation, consistent with the proposed mechanism of neural activity (and associated cognitive information) propagating across paths weighted by restFC. Another limitation to a reliance on data-driven optimization is the evidenced risk of overfitting to noise in the training set (Onoda et al., 2017; Teipel et al., 2017; Fountain-Zaragoza et al., 2019). Whilst the need to test the generalizability of connectivity-based predictive models across out-of-set samples and scanner sites also applies to activity flow, it is possible that an increased emphasis on neuroscientific theory (e.g. use of the healthy activation template rather than abstract coefficients) will not only ease translation of models to the clinic, but also quantitatively improve generalizability.

Nevertheless, it is worth highlighting that the reported activity flow results serve more as a proof-of-concept at this stage rather than a ready-to-implement clinical protocol. Future extensions specifically targeting improvement in the individualized (subject-level) prediction accuracy will be necessary to elevate variance explained to clinically useful thresholds. To this end, some of the optimization methods adopted in the more data-driven fingerprinting approaches cited above may be integrated with activity flow mapping (e.g. nested cross-validation of restFC estimation parameters to optimize prediction accuracy). Critically, such optimization is only advocated if it complements the model’s underlying theory. Indeed, the observed improvement in subject-level prediction accuracy with the two-cycle activity flow approach highlights potential for future optimization that is theoretically grounded (in this case, by similar transformations operationalized in neural network models). Collecting higher quality fMRI data (e.g. higher spatiotemporal resolution sequences, Smith et al., 2013), longer resting-state scan durations per subject (Gordon et al., 2017) and encompassing multiple scanner sites (Woo et al., 2017), all serve as important data acquisition refinements that may also improve individualized prediction accuracy. Overall, the present findings favor a synergistic approach, wherein the theoretical constraints imposed by the activity flow framework allow for careful optimization of predictive frameworks whilst mitigating the risk of overfitting, potentially enabling better generalization in large out-of-set populations.

In pursuing these extensions of activity flow mapping, it is worth reiterating the practical benefits of the approach: predictions of cognitive task activations (and associated dysfunction) are made without having subjects actually perform in-scanner tasks. The resulting task activation predictions show promise for use in a variety of clinical contexts: to expedite diagnosis, identify targets for stimulation-based intervention (Fox et al., 2014; Reinhart and Nguyen, 2019), and for longitudinal assessment of prognosis/response to intervention (He et al., 2007). The reported preliminary success of activity flow mapping hence provides a theoretical foundation for future developments in the use of restFC as a clinical biomarker, aiding the realization of long-upheld goals of incorporating functional neuroimaging measures in personalized medicine applications.

## Acknowledgements

The authors acknowledge support by the US National Institutes of Health under awards R01 AG055556 and R01 MH109520 to MWC, awards P50 AG005681 and P01 AG026276 that enabled data collection by the Adult Children Study at the Knight Alzheimer’s Disease Research Center at Washington University in St Louis (Knight ADRC), and the K01 AG053474 awarded to BAG. The content is solely the responsibility of the authors and does not necessarily represent the official views of any of the funding agencies. The authors would also like to thank John C. Morris and Tammie L.S. Benzinger for their involvement in the Knight ADRC project, which facilitated the present report. The authors acknowledge the Office of Advanced Research Computing (OARC) at Rutgers, The State University of New Jersey for providing access to the Amarel cluster and associated research computing resources that have contributed to the results reported here.

